# Conserved changes in secondary structure and aggregation properties of *in vitro* evolved proteins for thermo stability

**DOI:** 10.1101/111443

**Authors:** Satyamurthy Kundharapu

## Abstract

Most of the screening strategies of directed evolution involved in thermo stability deals with aggregation of proteins either directly or indirectly. Here in this work I investigated what happens in aggregation property and secondary structure of the protein when it improved its thermo stability by incorporating certain amino acid changes in the protein. To study these changes I picked randomly 12 different proteins and I analyzed their 25 different thermo stable mutants. I used open access online Software to get the aggregation propensity values and values for different secondary structure elements propensities of proteins. I compared the aggregation propensity and predicted secondary structure values of thermo stable mutants with their parent Wild type proteins. The stable mutants followed three different conserved patterns to improve their thermo stability.

## Introduction

There are different strategies employed to improve the thermo stability of the protein like modifications, oligomerisation, domain shuffling, protein engineering. Among these strategies protein engineering is widely used for its promising positive results for thermo stability improvement in proteins which are having industrial and therapeutic applications. Most of the screening strategies of protein engineering involved in thermo stability deals with aggregation of proteins either directly or indirectly because, most of the proteins upon thermal denaturation will undergo to an irreversible aggregation. There is an unbreakable link in between the aggregation property and thermal melting of the protein. When you change the amino acids in proteins, along with changes in aggregation propensity values there will be a definite or slight changes will happened in the secondary structure of the protein.

Here in this work I want to study is this changes are following any trend among the thermo stable mutants. So in this study I compared trends in different parameters either decreasing or increasing.

To study these changes I used Online Software to get the aggregation propensity values and predicted propensity values for different secondary structure elements of proteins. I compared the aggregation propensity and predicted secondary structure values of thermo stable mutants with their parent Wild type proteins which are less stable than mutants at given temperature. For this study, I picked randomly 12 different proteins and their 25 different thermo stable variants (5,6,7,8,9,10,11,12,14).

## Methods and Material

### Proteins Used in this Study

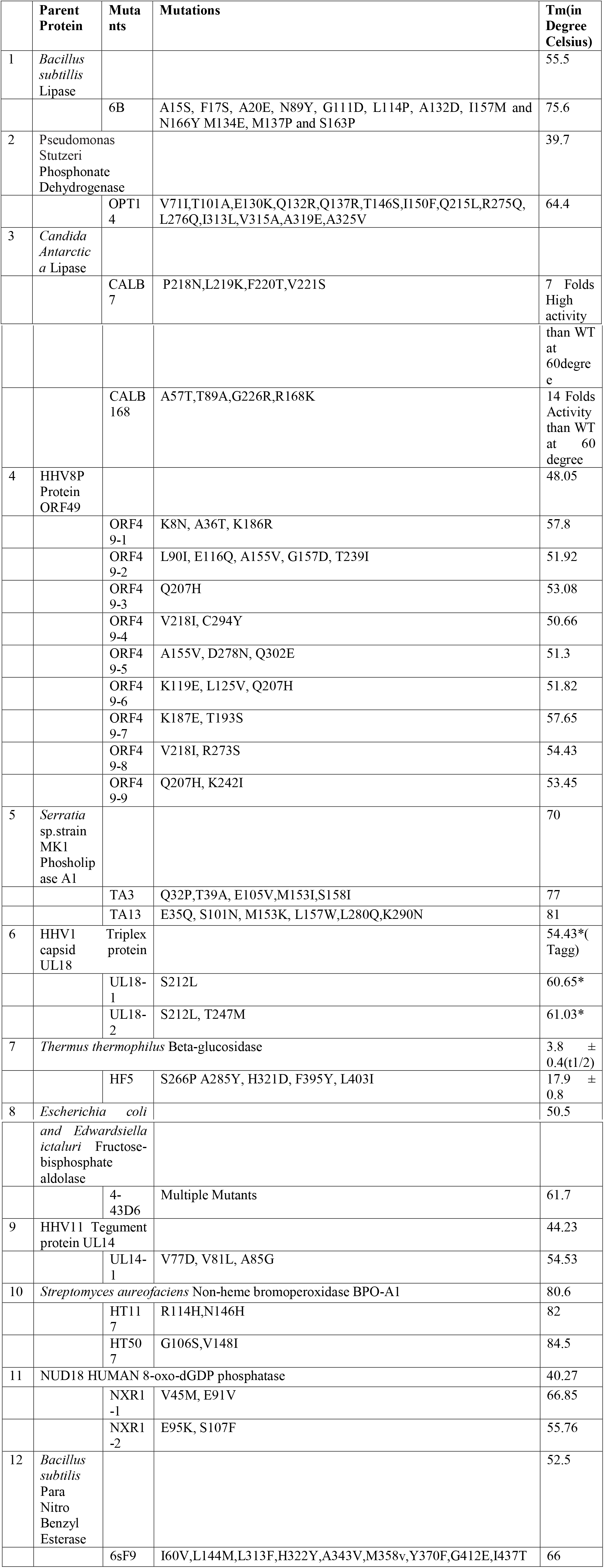

### Software Used in this Study

Four different software freely available online software were used in this study. Those are TANGO, AGGRESCAN, PASTA2.0 and CHOUFASMAN.

http://tango.crg.es/protected/academic/calculation.jsp

http://protein.bio.unipd.it/pasta2/

http://cho-fas.sourceforge.net/

http://bioinf.uab.es/aggrescan/

For total protein aggregation propensity values, TANGO Agg and Amylo values, AGGRESCAN a3vSA value and in PASTA2.0 number of Amyolids and for Secondary structure prediction TANGO Secondary structure propensities, PASTA Percentage of different secondary structure elements, CHOUFASMAN Percentage of different secondary structure elements were taken(1,2,3,4). Every software I used at their default settings. For analyzing the Local aggregation propensity value changes at the mutated regions I used TANGO (helix agg or Beta agg) and AGGRESCAN (a3v) (10, 12). To check local secondary structure values TANGO Helix propensity at the mutated region or Beta sheet propensity or Turn propensity and CHOUFASMAN were used. To check aggregation prone regions in the protein an open access tool WALTZ was used (Data not shown). For a mutant out of three different values from different software which two give a similar trend like either decrease or increase when compared with wild type that trend was considered for that mutant.

When there are three different software values for a given mutant, TANGO values of aggregation propensity and secondary structure prediction propensities were taken consideration or the changes in these parameters at that particular mutated region were analyzed.

## Results and discussion

### Lipase

**Table-1:** Predicted secondary structure propensities in WT and Thermo stable Mutant (6B)

|  | **WT** | **6B** |  | **WT** | **6B** |  | **WT** | **6B** |
| --- | --- | --- | --- | --- | --- | --- | --- | --- |
| **CFSSP Helix** | 55.2 | 49.2 | **PASTA Helix** | 27.62 | 20.99 | **TANGO Helix** | 37.4528 | 33.2371 |
| **CFSSP Beta** | 43.6 | 43.1 | **PASTA Beta** | 26.52 | 28.73 | **TANGO Beta** | 401.536 | 353.099 |
| **CFSSP Turn** | 10.5 | 11.6 | **PASTA Coil** | 45.86 | 50.28 | **Tango Turn** | 140.245 | 150.232 |

**Table-2.**
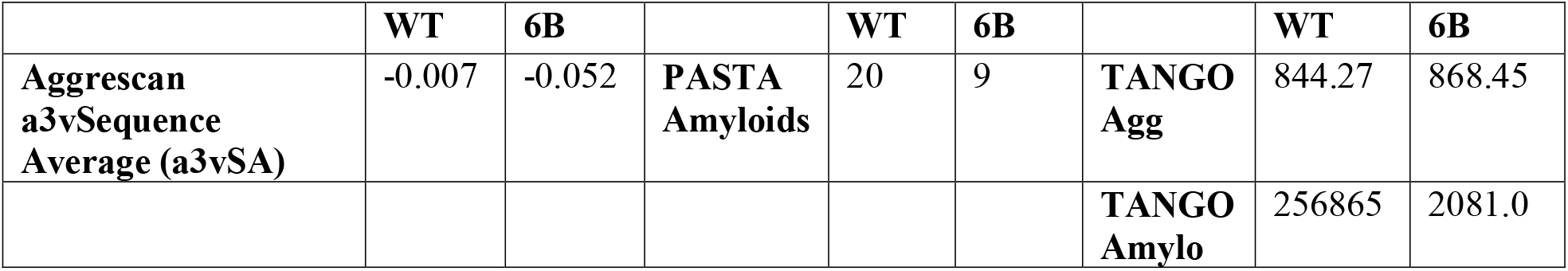
Predicted aggregation propensity values of WT and Thermo Stable Mutant (6B)

The comparison of predicted secondary structure elements propensities of wild type lipase with stabilized mutant 6B showed there is a decrease in alpha helical content and beta sheet in 6B but there is an increase in overall Turn content in 6B. Comparison of wild type aggregation propensity values with 6B showed significant decrease. There is slight increase in Tango agg value of 6B but Tango Amylo value is too less for 6B than wildtype. At the every mutated position the aggregation propensity values were decreased (data was not shown).This aggregation resistance behavior of 6B was reported earlier also (5).

### Phosphonate dehydrogenase

**Table-3.** Predicted secondary structure propensities in WT and Thermo stable Mutant (OPT14)

|  | WT | OPT14 |  | WT | OPT14 |  | WT | OPT14 |
| --- | --- | --- | --- | --- | --- | --- | --- | --- |
| **CFSSP Helix** | 85.4 | 84.5 | **PASTA Helix** | 54.46 | 57.74 | **Tango Helix** | 934.86 | 962.18 |
| **CFSSP Beta** | 38.7 | 36.3 | **PASTA Beta** | 8.03 | 6.25 | **Tango Beta** | 494.941 | 463.198 |
| **CFSSP Turn** | 9.2 | 9.2 | **PASTA Coil** | 37.5 | 36.01 | **Tango Turn** | 128.275 | 129.577 |

**Table-4.** Predicted aggregation propensity values of WT and Thermo Stable Mutant (OPT14)

|  | WT | OPT14 |  | WT | OPT14 |  | WT | OPT14 |
| --- | --- | --- | --- | --- | --- | --- | --- | --- |
| **PASTA Amyloids** | 6 | 4 | **a3vSA** | 0.005 | -0.001 | **Tango agg** | 1728.15 | 1556.57 |
|  |  |  |  |  |  | **Tango amylo** | 194968 | 170539 |

Opt14 (thermo stable mutant of phosphonate dehydrogenase) mutation Q215 to L showed increase in alpha helical content at 213-229 compared to wild type protein and the overall alpha helical content also improved. There is a significant decrease in overall aggregation propensity values and beta sheet values also observed in Opt 14 when compared with wild type.

### Candida Antarctica Lipase

**Table-5.** Predicted secondary structure propensities in WT and Thermo stable Mutants (CALB7, CALB168)

|  | WT | CAL B7 | CALB 168 |  | WT | CAL B7 | CALB 168 |  | WT | CAL B7 | CALB 168 |
| --- | --- | --- | --- | --- | --- | --- | --- | --- | --- | --- | --- |
| **PAST A Helix** | 22.7 | 23.4 | 25.87 | **CFSS P Helix** | 49.8 | 46.7 | 55.2 | **Tango Helix** | 110.9 59 | 110.9 56 | 82.284 4 |
| **aPAS TA Beta** | 13.25 | 10.73 | 11.36 | **CFSS P Beta** | 68.5 | 66.6 | 68.1 | **Tango Beta** | 683.4 25 | 671.4 82 | 687.80 5 |
| **PAST A Coil** | 64.4 | 65.93 | 62.78 | **CFSS P Turn** | 15.1 | 15.5 | 15.1 | **Tango Turn** | 182.7 55 | 186.4 46 | 178.49 2 |

**Table-6.**
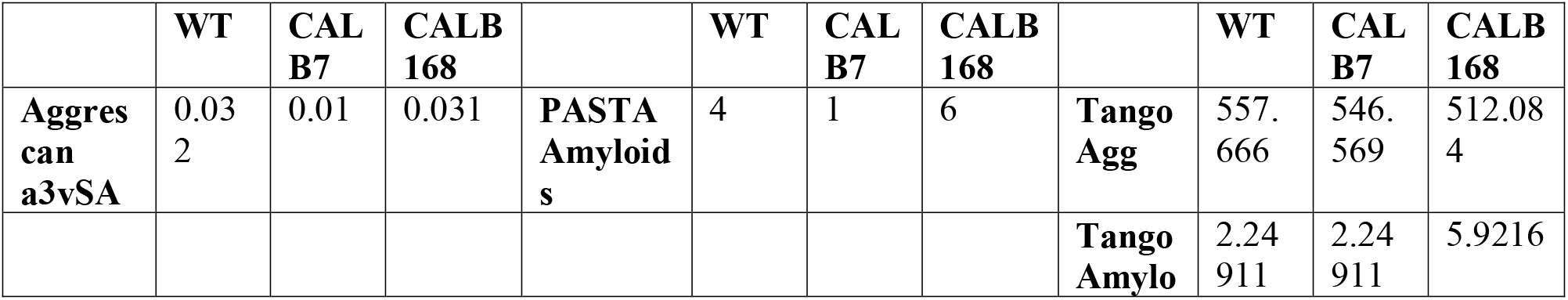
Predicted aggregation propensity values of WT and Thermo Stable Mutants (CALB7, CALB168)

Candida Antarctica lipase thermo stable mutants CALB7 and CALB168 stabilized in two different ways.CALB7 stabilized by decreasing the aggregation Propensity and CALB168 improved it’s stability by improving Alpha helix content and there is slight decrease in beta sheet also observed.

## ORF49

**Table-7.** Predicted secondary structure propensities in WT and Thermo stable Mutants By Choufasman algorithm

|  | WT | ORF4<br>9-1 | ORF4<br>9-2 | ORF4<br>9-3 | ORF4<br>9-4 | ORF4<br>9-5 | ORF4<br>9-6 | ORF4<br>9-7 | ORF4<br>9-8 | ORF4<br>9-9 |
| --- | --- | --- | --- | --- | --- | --- | --- | --- | --- | --- |
| **CFSSP**<br>**Helix** | 74.5 | 73.8 | 75.5 | 74.5 | 74.5 | 74.5 | 74.5 | 74.5 | 74.5 | 74.5 |
| **CFSSP**<br>**Beta** | 82.5 | 82.5 | 82.8 | 82.5 | 82.5 | 83.1 | 82.5 | 82.5 | 82.5 | 82.5 |
| **CFSSP**<br>**Turn** | 8.6 | 8.6 | 8.6 | 8.6 | 8.6 | 8.6 | 8.6 | 8.6 | 8.6 | 8.3 |

**Table-8.** Predicted secondary structure propensities in WT and Thermo stable Mutants by PASTA2.0

|  | WT | ORF4<br>9-1 | ORF4<br>9-2 | ORF4<br>9-3 | ORF4<br>9-4 | ORF4<br>9-5 | ORF4<br>9-6 | ORF4<br>9-7 | ORF4<br>9-8 | ORF4<br>9-9 |
| --- | --- | --- | --- | --- | --- | --- | --- | --- | --- | --- |
| **PASTA**<br>**Helix** | 72.19 | 66.87 | 74.83 | 72.52 | 72.52 | 72.19 | 70.53 | 73.84 | 72.19 | 72.52 |
| **PASTA**<br>**Beta** | 0.66 | 0.99 | 0.33 | 0.66 | 0.66 | 0.99 | 1.32 | 0.33 | 0.66 | 0.66 |
| **PASTA**<br>**Coil** | 27.15 | 30.13 | 24.83 | 26.82 | 26.82 | 26.82 | 28.15 | 25.83 | 27.15 | 26.82 |

**Table-9.** Predicted secondary structure propensities in WT and Thermo stable Mutants by TANGO

|  | WT | ORF4<br>9-1 | ORF4<br>9-2 | ORF4<br>9-3 | ORF4<br>9-4 | ORF4<br>9-5 | ORF4<br>9-6 | ORF4<br>9-7 | ORF4<br>9-8 | ORF4<br>9-9 |
| --- | --- | --- | --- | --- | --- | --- | --- | --- | --- | --- |
| **Tango**<br>**Helix** | 138.41<br>6 | 129.25<br>9 | 161.37<br>4 | 137.61<br>6 | 141.32<br>8 | 142.15<br>7 | 121.09<br>2 | 137.45 | 138.17<br>3 | 139.27<br>9 |
| **Tango**<br>**Beta** | 757.46<br>8 | 769.18<br>9 | 751.53<br>2 | 755.92<br>9 | 751.54<br>5 | 762.23<br>7 | 771.32<br>9 | 756.55<br>2 | 746.49<br>8 | 758.71 |
| **Tango**<br>**Turn** | 138.64<br>7 | 138.90<br>5 | 136.86<br>8 | 139.42<br>2 | 138.29<br>6 | 132.94<br>9 | 138.56<br>4 | 138.79 | 138.68<br>7 | 139.00<br>6 |

**Table-10.** Predicted aggregation propensity values of WT and Thermo Stable Mutants by Tango

|  | WT | ORF4<br>9-1 | ORF4<br>9-2 | ORF4<br>9-3 | ORF4<br>9-4 | ORF4<br>9-5 | ORF4<br>9-6 | ORF4<br>9-7 | ORF4<br>9-8 | ORF4<br>9-9 |
| --- | --- | --- | --- | --- | --- | --- | --- | --- | --- | --- |
| **Tango Agg** | 2752.09 | 2657.27 | 2778.93 | 2720.23 | 2751.31 | 2730.21 | 2721.36 | 2483.02 | 2758 | 3191.66 |
| **Tango Amylo** | 275053 | 161947 | 275054 | 275053 | 275053 | 275054 | 275054 | 275054 | 275054 | 275054 |

**Table-11.** Predicted aggregation propensity values of WT and Thermo Stable Mutants by Aggrescan

|  | **WT** | **ORF4 9-1** | **ORF4 9-2** | **ORF4 9-3** | **ORF4 9-4** | **ORF4 9-5** | **ORF4 9-6** | **ORF4 9-7** | **ORF4 9-8** | **ORF4 9-9** |
| --- | --- | --- | --- | --- | --- | --- | --- | --- | --- | --- |
| **Aggrescan a3vSA** | 0.089 | 0.086 | 0.092 | 0.09 | 0.092 | 0.095 | 0.092 | 0.087 | 0.093 | 0.099 |

**Table-12.** Predicted aggregation propensity values of WT and Thermo Stable Mutants by PASTA

|  | **WT** | **ORF4 9-1** | **ORF4 9-2** | **ORF4 9-3** | **ORF4 9-4** | **ORF4 9-5** | **ORF4 9-6** | **ORF4 9-7** | **ORF4 9-8** | **ORF4 9-9** |
| --- | --- | --- | --- | --- | --- | --- | --- | --- | --- | --- |
| **PASTA Amyloids** | 20 | 20 | 20 | 20 | 20 | 20 | 20 | 20 | 20 | 20 |

Thermo stability screening of ORF49 mutants is aggregation dependent, out of nine mutants ORF49-1, ORF49-7 showed decreased aggregation propensity values. ORF49-2 showed increase in alpha helical content. ORF49-3 (Q207H) a single mutant showed both improvement in alpha helix and decrease in aggregation propensity but ORF49-6 mutant had along with ORF49-3 mutation two extra mutations which are decreasing Tm by 2degrees of single mutant.ORF49-6 showing same aggregation propensity values like ORF49-3 but there is a significant decrease in alpha Helix content. The Decrease in Tm of ORF49-6 is due decrease in alpha helix content.ORF49-9 mutant also had this single mutation (Q207H) along with one more mutation and it doesn’t showed any significant change in Tm but there is increase in aggregation propensity values. Alpha Helix content of ORF49-9 is slightly more than ORF49-3. ORF49-4 and ORF49-8 had one common mutation (V218I) but there is Tm difference of 4 degrees present in between these mutants the second mutation in ORF49-4 improving the beta sheet, in ORF49-8 the second mutation causing slight increase in local alpha helical content. Previous mutations in different proteins showed increase beta sheet content decrease the stability so the decrease in Tm of ORF49-4 is due increase in Beta sheet. ORF49-5 showed improvement in alpha helix content and showed decrease in aggregation propensity value.

### Phospholipase A1

**Table-13.** Predicted secondary structure propensities in WT and Thermo stable Mutants (TA3,TA13)

|  | **WT** | **TA3** | **TA13** |  | **WT** | **TA3** | **TA13** |  | **WT** | **TA3** | **TA13** |
| --- | --- | --- | --- | --- | --- | --- | --- | --- | --- | --- | --- |
| **CFSS P Helix** | 67.8 | 69.1 | 67.2 | **PAST A Helix** | 44.06 | 44.06 | 43.13 | **Tango Helix** | 302.898 | 324.614 | 294.616 |
| **CFSS P Beta** | 60.3 | 61.16 | 60.3 | **PAST A Beta** | 8.13 | 9.38 | 8.44 | **Tango Beta** | 601.823 | 626.761 | 591.179 |
| **CFSS**<br>**P**<br>**Turn** | 12.5 | 12.8 | 13.1 | **PAST**<br>**A Coil** | 47.18 | 46.56 | 48.44 | **Tango**<br>**Turn** | 123.8<br>58 | 116.4<br>39 | 128.0<br>95 |

**Table-14.** Predicted aggregation propensity values of WT and Thermo Stable Mutants (TA3, TA13)

|  | **WT** | **TA3** | **TA13** |  | **W**<br>**T** | **TA**<br>**3** | **TA1**<br>**3** |  | **WT** | **TA3** | **TA13** |
| --- | --- | --- | --- | --- | --- | --- | --- | --- | --- | --- | --- |
| **Aggresc**<br>**an**<br>**a3vSA** | -<br>0.104 | 2.81<br>2 | -<br>0.122 | **PASTA**<br>**Amyloi**<br>**ds** | 0 | 2 | 0 | **Tango**<br>**Agg** | 1427<br>.32 | 1469.8<br>6 | 1428.4<br>2 |
|  |  |  |  |  |  |  |  | **Tango**<br>**Amylo** | 4016<br>32 | 35208<br>3 | 39130<br>1 |

In Phospholipase mutants TA13 improved its aggregation resistance. It was also observed that, mutated regions had less aggregation propensity values than WT protein. TA3 doesn’t show any decrease in aggregation propensity values but improved its alpha helical content.

### Triplex capsid protein UL18

**Table-15.** Predicted secondary structure propensities in WT and Thermo stable Mutants (UL18-1, UL18-2)

|  | **WT** | **UL18**<br>**-1** | **UL18**<br>**-2** |  | **WT** | **UL18**<br>**-1** | **UL18**<br>**-2** |  | **WT** | **UL18-**<br>**1** | **UL18-**<br>**2** |
| --- | --- | --- | --- | --- | --- | --- | --- | --- | --- | --- | --- |
| **CFSS**<br>**P**<br>**Helix** | 70.1 | 71.7 | 71.7 | **PAST**<br>**A**<br>**Helix** | 35.22 | 35.22 | 35.22 | **Tang**<br>**o**<br>**Helix** | 809.33<br>1 | 809.71<br>8 | 800.11<br>4 |
| **CFSS**<br>**P**<br>**Beta** | 47.2 | 47.2 | 47.2 | **PAST**<br>**A Beta** | 16.98 | 16.98 | 16.98 | **Tang**<br>**o**<br>**Beta** | 687.58<br>5 | 671.07<br>1 | 663.69<br>5 |
| **CFSS**<br>**P**<br>**Turn** | 11.3 | 11.3 | 11 | **PAST**<br>**A coil** | 47.8 | 47.8 | 47.8 | **Tang**<br>**o**<br>**Turn** | 140.64<br>3 | 140.81<br>6 | 140.8 |

**Table-16.** Predicted aggregation propensity values of WT and Thermo Stable Mutants (UL18-1, UL18-2)

|  | **WT** | **UL1**<br>**8-1** | **UL1**<br>**8-2** |  | **WT** | **UL1**<br>**8-1** | **UL1**<br>**8-2** |  | **WT** | **UL1**<br>**8-1** | **UL1**<br>**8-2** |
| --- | --- | --- | --- | --- | --- | --- | --- | --- | --- | --- | --- |
| **PAST**<br>**Amyloi**<br>**ds** | 20 | 20 | 20 | **Aggresc**<br>**an**<br>**a3vSA** | 0.09 | 0.095 | 0.099 | **Tango**<br>**Agg** | 1846.<br>89 | 2008.<br>86 | 2008.<br>89 |
|  |  |  |  |  |  |  |  | **Tango**<br>**Amylo** | 20.36<br>24 | 20.36<br>24 | 20.36<br>24 |

Both thermo stable mutants of UL18 had common mutation (S212L) which increases the Local Alpha helix content decreases the beet sheet values and the UL18-2 had one more mutation that decreases the beet sheet value furthermore. The decrease in beta sheet value and local improvement alpha helix is cause of stabilization both mutants.

### Beta-glucosidase Thermus thermophiles

**Table-17.** Predicted secondary structure propensities in WT and Thermo stable Mutant (HF5)

|  | WT | HF5 |  | WT | HF5 |  | WT | HF5 |
| --- | --- | --- | --- | --- | --- | --- | --- | --- |
| **CFSSP Helix** | 63.8 | 62.6 | **PASTA Helix** | 45.24 | 44.78 | **Tango Helix** | 772.383 | 899.635 |
| **CFSSP Beta** | 57.1 | 57.1 | **PASTA Beta** | 9.74 | 10.21 | **Tango Beta** | 757.952 | 747.42 |
| **CFSSP Turn** | 12.8 | 12.8 | **PASTA Coil** | 45.01 | 45.01 | **Tango Turn** | 139.408 | 134.842 |

**Table-18.** Predicted aggregation propensity values of WT and Thermo Stable Mutant (HF5)

|  | WT | HF5 |  | WT | HF5 |  | WT | HF5 |
| --- | --- | --- | --- | --- | --- | --- | --- | --- |
| **PASTA Amyloids** | 7 | 7 | **Aggrescan a3vSA** | -0.014 | -0.013 | **Tango Agg** | 1971.09 | 2053 |
|  |  |  |  |  |  | **Tango Amylo** | 1.35E+06 | 1.35E+06 |

Thermo stable mutant of Beta glucosidase showed significant increase in alpha helical values at the mutated regions. Showed decrease beta sheet values. But there is no significant change in overall aggregation propensity values although two mutations of this protein showed little decrease at their positions.

### Fructose bisphosphate aldolase

**Table-19.** Predicted secondary structure propensities in WT and Thermo stable Mutant (4-43D6)

|  | WT | 4-43D6 |  | WT | 4-43D6 |  | WT | 4-43D6 |
| --- | --- | --- | --- | --- | --- | --- | --- | --- |
| **CFSSP Helix** | 72.6 | 73.1 | **PASTA Helix** | 43.58 | 43.7 | **Tango Helix** | 165.262 | 266.39 |
| **CFSSP Bete** | 32.4 | 41.7 | **PASTA Beta** | 14.25 | 14.85 | **Tango Beta** | 854.948 | 817.164 |
| **CFSSP Turn** | 13.4 | 12.3 | **PASTA Coil** | 42.18 | 41.46 | **Tango Turn** | 154.457 | 152.639 |

**Table-20.**
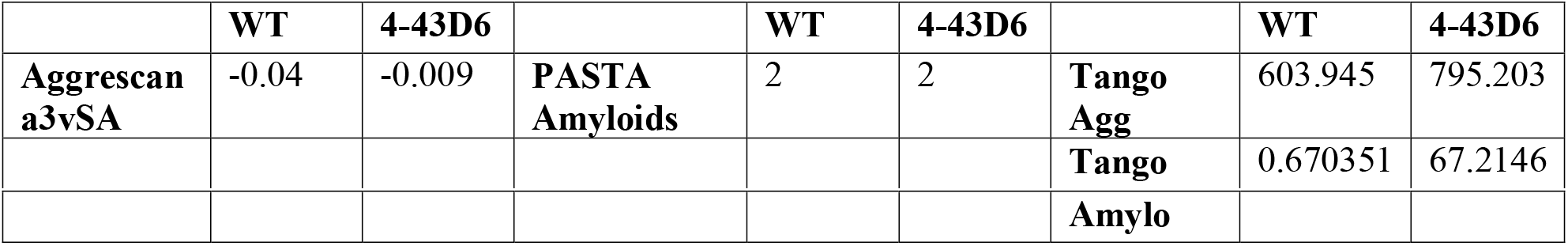
Predicted aggregation propensity values of WT and Thermo Stable Mutant (4-43D6)

Aggregation propensity values for WT protein is too less. Thermo stable mutant 4-43D6 showed increase alpha helix content and also showed there is an increase aggregation propensity values. But this mutant stabilized because of improvement in alpha helix content.

### Tegument UL14

**Table-21.** Predicted secondary structure propensities in WT and Thermo stable Mutants (UL14-1)

|  | WT | UL14-1 |  | WT | UL14-1 |  | WT | UL14-1 |
| --- | --- | --- | --- | --- | --- | --- | --- | --- |
| **CFSSP Helix** | 67.6 | 67.6 | **PASTA Helix** | 52.97 | 53.88 | **Tango Helix** | 523.461 | 523.489 |
| **CFSSP Beta** | 36.1 | 36.1 | **PASTA Beta** | 1.83 | 0 | **Tango Beta** | 312.36 | 272.687 |
| **CFSSP Turn** | 15.5 | 16 | **PASTA Coil** | 45.21 | 46.12 | **Tango Turn** | 125.093 | 126.852 |

**Table-22.** Predicted aggregation propensity values of WT and Thermo Stable Mutant (UL14-1)

|  | WT | UL14-1 |  | WT | UL14-1 |  | WT | UL14-1 |
| --- | --- | --- | --- | --- | --- | --- | --- | --- |
| **PASTA Amyloids** | 0 | 0 | **Aggrescan a3vSA** | -0.266 | -0.285 | **Tango Agg** | 287.803 | 287.803 |
|  |  |  |  |  |  | **Tango Amylo** | 0.994051 | 0.994045 |

Aggregation propensity values for WT protein is too less. Aggrescan values showed there is an aggregation resistance although there is change in Tango values. Thermo stable mutant of this viral protein showed significant increase in alpha Helix content.

### Non-heme bromoperoxidase

**Table-23.** Predicted secondary structure propensities in WT and Thermo stable Mutants (HT117, HT507)

|  | WT | HT117 | HT507 |  | WT | HT117 | HT507 |  | WT | HT117 | HT507 |
| --- | --- | --- | --- | --- | --- | --- | --- | --- | --- | --- | --- |
| **CFSSP Helix** | 68 | 68 | 69.5 | **PASTA Helix** | 35.27 | 35.64 | 32 | **Tango Helix** | 181.586 | 159.313 | 186.417 |
| **CFSSP Beta** | 33.8 | 33.8 | 33.8 | **PASTA Beta** | 19.64 | 19.64 | 21.82 | **Tango Beta** | 478.027 | 477.883 | 483.261 |
| **CFSSP Turn** | 13.5 | 13.1 | 13.5 | **PASTA Coil** | 45.09 | 44.73 | 46.18 | **Tango Turn** | 160.112 | 149.923 | 166.113 |

**Table-24.** Predicted aggregation propensity values of WT and Thermo Stable Mutant (HT117, HT507)

|  | WT | HT117 | HT507 |  | WT | HT117 | HT507 |  | WT | HT117 | HT507 |
| --- | --- | --- | --- | --- | --- | --- | --- | --- | --- | --- | --- |
| **PASTA Amyloids** | 4 | 5 | 4 | **Aggrecan a3vSA** | - 0.103 | - 0.102 | - 0.106 | **Tango Agg** | 740.8 37 | 761.5 13 | 741.1 29 |
|  |  |  |  |  |  |  |  | **Tango Amylo** | 5716 7.2 | 5716 7.6 | 5693 3.9 |

The mutant HT117 mutation in the position 114 showed decrease beta sheet value but second mutation showed increase in beta sheet value and decrease in alpha helix value. But second thermo stable mutant HT507 showed significant decrease in aggregation propensity values and increase in alpha helix values. In Comparison both mutants HT507 is more thermo stable than HT117 although the mutations located same regions but different positions.

### NUD18 HUMAN 8-oxo-dGDP phosphatase

**Table-25.** Predicted secondary structure propensities in WT and Thermo stable Mutants (NXR1-1, NXR1-2)

|  | WT | NXR 1-1 | NXR 1-2 |  | WT | NXR 1-1 | NXR 1-2 |  | WT | NXR 1-1 | NXR 1-2 |
| --- | --- | --- | --- | --- | --- | --- | --- | --- | --- | --- | --- |
| **CFSS P Helix** | 71.2 | 71.2 | 71.2 | **PAST A Helix** | 42.14 | 43.65 | 41.18 | **Tango Helix** | 475.3 49 | 485.1 76 | 544.6 06 |
| **CFSS P Beta** | 48 | 49.2 | 48 | **PAST A Beta** | 13.13 | 12.38 | 14.24 | **Tango Beta** | 762.0 48 | 719.8 33 | 748.5 21 |
| **CFSS P Turn** | 12.7 | 12.4 | 12.7 | **PAST A Coil** | 44.27 | 43.96 | 44.58 | **Tango Turn** | 87.18 18 | 87.13 66 | 87.35 29 |

**Table-26.** Predicted aggregation propensity values of WT and Thermo Stable Mutants (NXR1-1, NXR1-2)

|  | WT | NXR 1-1 | NXR 1-2 |  | WT | NXR 1-1 | NXR 1-2 |  | WT | NXR 1-1 | NXR 1-2 |
| --- | --- | --- | --- | --- | --- | --- | --- | --- | --- | --- | --- |
| **Aggrecan a3vSA** | 0.006 | 0.013 | 0.014 | **PASTA Amyloids** | 20 | 20 | 20 | **Tango Agg** | 3191. 53 | 3192. 64 | 3225. 56 |
|  |  |  |  |  |  |  |  | **Tango Amylo** | 3165 3.7 | 5597 7.9 | 3165 2.9 |

The First thermo stable mutant NXR1-1 showed significant increase in Alpha helix. There is decrease in beta sheet compared to second mutant NXR1-2. NXR1-2 also showed significant increase in alpha helix at their mutated regions and decrease in beta sheet value. The aggregation propensity values when compared with WT slightly increased.

### p-Nitro Benzyl Esterase

**Table-27.** Predicted secondary structure propensities in WT and Thermo stable Mutant (6sF9)

|  | WT | 6SF9 |  | WT | 6SF9 |  | WT | 6SF9 |
| --- | --- | --- | --- | --- | --- | --- | --- | --- |
| **CFSSP Helix** | 76.5 | 76.3 | **PASTA Helix** | 40.7 | 41.2 | **Tango Helix** | 793.101 | 1015.04 |
| **CFSSP Beta** | 28.8 | 29.7 | **PASTA Beta** | 15.54 | 14.72 | **Tango Beta** | 1099.84 | 1085.02 |
| **CFSSSP Turn** | 14.5 | 14.5 | **PASTA Coil** | 43.76 | 43.56 | **Tango Turn** | 272.585 | 256.468 |

**Table-28.** Predicted aggregation propensity values of WT and Thermo Stable Mutant (6sF9)

|  | WT | 6SF9 |  | WT | 6SF9 |  | WT | 6SF9 |
| --- | --- | --- | --- | --- | --- | --- | --- | --- |
| **Aggrescan a3vSA** | -0.035 | -0.037 | **PASTA Amyloids** | 20 | 20 | **Tango Agg** | 2346.69 | 2277.06 |
|  |  |  |  |  |  | **Tango Amylo** | 3626.39 | 32.9687 |

p-Nitro Benzyl Esterase thermo stable mutant 6sF9 showed an improvement in overall alpha helix content and decrease in overall aggregation propensity values and there is a decrease in beta sheet value also.

## Conclusion

All thermo stable mutants analyzed in this study, out of 25 mutants 18 showed either there is decrease in aggregation propensity values or increase in alpha helix secondary structure prediction values. Remaining 7 mutants showed these changes at their mutated regions and almost of all of this mutants showed the decrease in overall beta sheet values and it was already reported that involvement beta sheet in thermal aggregation (13). The proteins which are thermo stabilized analyzed in this study showed are tabulated into three different groups.

**Table-29.**
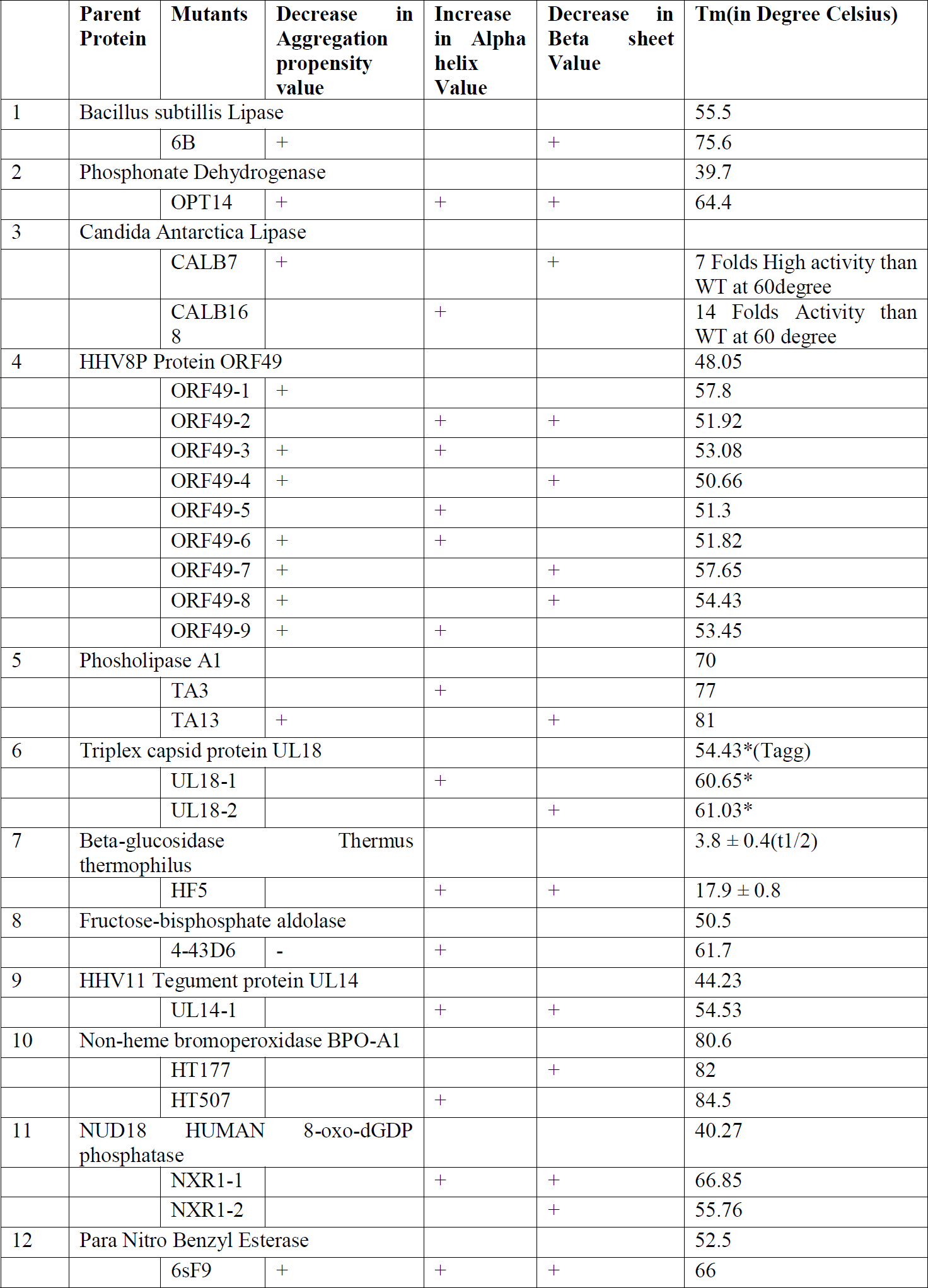
Changes in three different parameters happened in stabilized mutants

|  | Parent Protein | Mutants | Decrease in Aggregation propensity value | Increase in Alpha helix Value | Decrease in Beta sheet Value | Tm(in Degree Celsius) |
| --- | --- | --- | --- | --- | --- | --- |
| 1 | Bacillus subtilis Lipase |  |  |  |  | 55.5 |
|  |  | 6B | + |  | + | 75.6 |
| 2 | Phosphonate Dehydrogenase |  |  |  |  | 39.7 |
|  |  | OPT14 | + | + | + | 64.4 |
| 3 | Candida Antarctica Lipase |  |  |  |  |  |
|  |  | CALB7 | + |  | + | 7 Folds High activity than WT at 60degree |
|  |  | CALB168 |  | + |  | 14 Folds Activity than WT at 60 degree |
| 4 | HHV8P Protein ORF49 |  |  |  |  | 48.05 |
|  |  | ORF49-1 | + |  |  | 57.8 |
|  |  | ORF49-2 |  | + | + | 51.92 |
|  |  | ORF49-3 | + | + |  | 53.08 |
|  |  | ORF49-4 | + |  | + | 50.66 |
|  |  | ORF49-5 |  | + |  | 51.3 |
|  |  | ORF49-6 | + | + |  | 51.82 |
|  |  | ORF49-7 | + |  | + | 57.65 |
|  |  | ORF49-8 | + |  | + | 54.43 |
|  |  | ORF49-9 | + | + |  | 53.45 |
| 5 | Phosholipase A1 |  |  |  |  | 70 |
|  |  | TA3 |  | + |  | 77 |
|  |  | TA13 | + |  | + | 81 |
| 6 | Triplex capsid protein UL18 |  |  |  |  | 54.43*(Tagg) |
|  |  | UL18-1 |  | + |  | 60.65* |
|  |  | UL18-2 |  |  | + | 61.03* |
| 7 | Beta-glucosidase Thermus thermophilus |  |  |  |  | 3.8 ± 0.4(t1/2) |
|  |  | HF5 |  | + | + | 17.9 ± 0.8 |
| 8 | Fructose-bisphosphate aldolase |  |  |  |  | 50.5 |
|  |  | 4-43D6 | - | + |  | 61.7 |
| 9 | HHV11 Tegument protein UL14 |  |  |  |  | 44.23 |
|  |  | UL14-1 |  | + | + | 54.53 |
| 10 | Non-heme bromoperoxidase BPO-A1 |  |  |  |  | 80.6 |
|  |  | HT177 |  |  | + | 82 |
|  |  | HT507 |  | + |  | 84.5 |
| 11 | NUD18 HUMAN 8-oxo-dGDP phosphatase |  |  |  |  | 40.27 |
|  |  | NXR1-1 |  | + | + | 66.85 |
|  |  | NXR1-2 |  |  | + | 55.76 |
| 12 | Para Nitro Benzyl Esterase |  |  |  |  | 52.5 |
|  |  | 6sF9 | + | + | + | 66 |

OPT14 (thermo stable mutant of phosphite dehydrogenase) and 6sF9 (thermo stable mutant of parnitro benzyl esterase) improved their Tm in a greater extent (OPT 25degrees, 6sF9 14.5degrees than their respective parent protein) with less number mutations. These thermo stable mutants of phosphite dehydrogenase and para nitro benzyl esterase showed all three existing parameters changed and this changes shown additive effect to improve the thermo stability of these mutants in greater extent. This *in vitro* evolved proteins for thermo stability showed three types conserved changes, including their secondary structure and aggregation propensity. Sometime this changes are local at the mutated region and doesn’t show global changes in protein structure, but others showed significant change at local mutated region and overall global change in the protein.

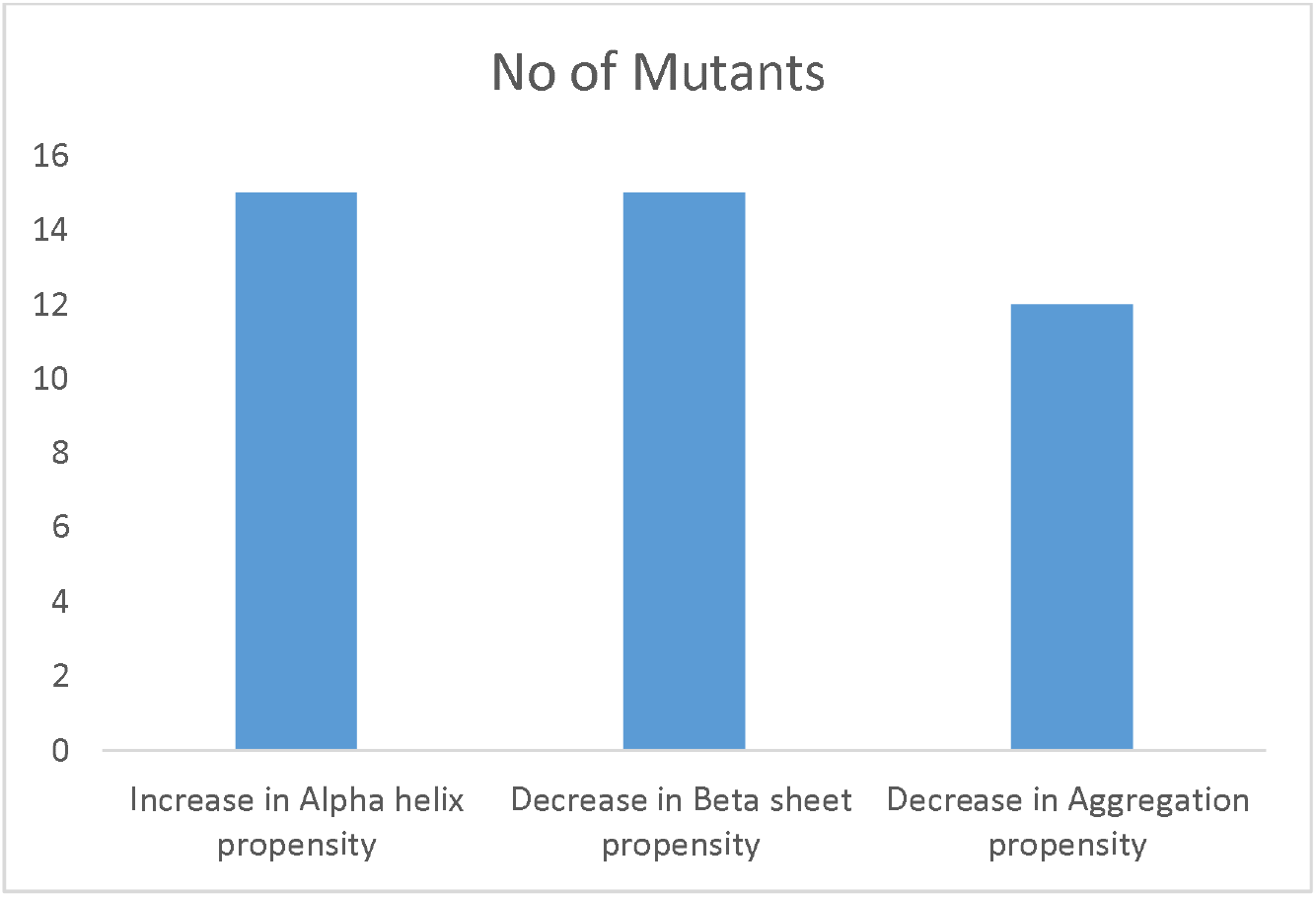
Changes in three different parameters happened in stabilized mutants

## Future Directions

This study can help in rational designing of protein for improving their stability in future to minimize the errors in it.

## References

1. Chou PY, Fasman GD (1974). “Conformational parameters for amino acids in helical, beta-sheet, and random coil regions calculated from proteins”. Biochemistry 13: 211–222.

2. Fernadez-Escamilla AM, Rousseau F, Schymkowtiz J, Serrano L, Prediction of sequence dependent mutational effects on the aggregation of peptide and protein, Nat Biotechnology, e-pub,2004.

3. Ian Walsh, Flavio Seno, Silvio C.E. Tosatto and Antonio Trovato. PASTA2: An improved server for protein aggregation prediction. Nucleic Acids Research, **accepted**. (2014)

4. Ignacio Asial, Yue Xiang Cheng, Henrik Engman, Binghuang Wu, Par̈ Nordlund & Tobias Cornvik, Engineering protein thermostability using a generic activity-independent biophysical screen Inside the cell. Nature Communications, Dec 2013.

5. Jae kwang song, Joon shick Rhee, Simultaneous enhancement of thermostability and catalytic activity of phospholipase A1 by evolutionary Molecular engineering. Applied Environmental Microbiology, Mar.2000, p.890–894

6. Jijun Hao and Alan Berry, A thermostable variant of fructose bisphosphate aldolase constructed by directed evolution also shows increased stability in organic solvents. Protein Engineering, Design & Selection vol. 17 no. 9 pp. 689–697, 2004

7. Kamal MZ, Ahmad S, Molugu TR, Vijayalakshmi A, Deshmukh MV, Sankaranarayanan R, Rao NM In vitro evolved non-aggregating and thermostable lipase: structural and thermodynamic investigation. J Mol Biol, 2011, 413:726–741.

8. Lori Giver, Anne Gershenson, Directed evolution of thermo stable Esterase. Proc. Natl. Acad. Sci, 1998, Vol95, 12809–12813.

9. Michael J. McLachlan, Future improvement of phosphite dehydrogenase thermo stability by saturation mutagenesis. Biotechnology bioengineering, 2008;99: 268–274.

10. Oscar Conchillo-Solé, Natalia S de Groot, Francesc X Avilés, Josep Vendrell,Xavier Daura and Salvador Ventura, AGGRESCAN: a server for the prediction and evaluation of “hot spots” of aggregation in polypeptides. BMC Bioinformatics 2007, 8:65

11. Ryosuke Yamada, Tatsutoshi Higo, Chisa Yoshikawa, Hideyasu China, Hiroyasu Ogino, Improvement of the stability and activity of the BPO-A1haloperoxidase from Streptomyces aureofaciens by directed evolution Journal of Biotechnology 192, 2014, 248–254

12. Xiang-Qian Peng, Improved Thermostability of Lipase B from Candida Antarctica by Directed Evolution and Display on Yeast Surface. Appl Biochem Biotechnol, 2013, 169:351–358

13. Yong-Bin Yan, Qi Wang, Hua-Wei He, and Hai-Meng Zhou* Protein Thermal Aggregation Involves Distinct Regions: Sequential Events in the Heat-Induced Unfolding and Aggregation of Hemoglobin Biophys J. 2004 Mar; 86(3): 1682–1690.

14. Zhuo-Lin Yi, Shuai-Bing Zhang a, Xiao-Qiong Pei, Zhong-Liu Wu, Design of mutants for enhanced thermostability of b-glycosidase BglY from Thermus thermophiles. Bioresource Technology 129, 2013, 629–633

